# Hippocampal atrophy and intrinsic brain network dysfunction relate to alterations in mind wandering in neurodegeneration

**DOI:** 10.1101/194092

**Authors:** Claire O’Callaghan, James M. Shine, John R. Hodges, Jessica R. Andrews-Hanna, Muireann Irish

**Author notes:** **Corresponding authors** Associate Professor Muireann Irish, Brain and Mind Centre and School of Psychology, the University of Sydney, 100 Mallet Street, Camperdown, NSW 2050, Australia. Claire O’Callaghan, Herchel Smith Building for Brain & Mind Sciences, Robinson Way, Cambridge CB2 0SZ, UK.

## Abstract

Mind wandering represents the human capacity for internally focussed thought, and relies upon the brain’s default network and its interactions with attentional networks. Studies have characterised mind wandering in healthy people, yet there is limited understanding of how this capacity is affected in clinical populations. This study used a validated thought-sampling task to probe mind wandering capacity in two neurodegenerative disorders: behavioural variant frontotemporal dementia (bvFTD; n=35) and Alzheimer’s disease (AD; n=24), compared to older controls (n=37). These patient groups were selected due to canonical structural and functional changes across sites of the default and frontoparietal networks, and well-defined impairments in cognitive processes that support mind wandering. Relative to controls, bvFTD patients displayed significantly reduced mind wandering capacity, offset by a significant increase in stimulus-bound thought. In contrast, AD patients demonstrated comparable levels of mind wandering to controls, in the context of a relatively subtle shift towards stimulus-/task-related forms of thought. In the patient groups, mind wandering was associated with grey matter integrity in the hippocampus/parahippocampus, striatum, insula and orbitofrontal cortex. Resting state functional connectivity revealed associations between mind wandering capacity and connectivity within and between regions of the frontoparietal and default networks, with distinct patterns evident in patients vs. controls. These findings support a relationship between altered mind wandering capacity in neurodegenerative disorders, and structural and functional integrity of the default and frontoparietal networks. This study highlights a dimension of cognitive dysfunction not well documented in neurodegenerative disorders, and validates current models of mind wandering in a clinical population.

**Significance statement:** Humans spend much of their waking life engaged in mind wandering. Underlying brain systems supporting this complex ability have been established in healthy individuals, yet it remains unclear how mind wandering is altered in neuropsychiatric populations. We reveal changes in the thought profiles elicited during periods of low cognitive demand in dementia, resulting in reduced mind wandering and an increased propensity towards stimulus-bound thought. These altered thought profiles were associated with structural and functional brain changes in the hippocampus, default and frontoparietal networks; key regions implicated in internal mentation in healthy individuals. Our findings provide a unique clinical validation of current theoretical models of mind wandering, and reveal a dimension of cognitive dysfunction that has received scant attention in dementia.

---

Mind wandering is fundamental to the human experience, yet its alteration in clinical populations remains poorly understood. Dynamic interactions within and between large-scale brain networks govern the initiation and maintenance of mind wandering (1, 2). Of particular interest in this context are interactions between the default network and the frontoparietal control network (3-5). In a recently proposed framework, spontaneous and unconstrained internally oriented thought is generated by fluctuations in the medial temporal lobe system of the default network, with weak influence from frontoparietal regions (2). More deliberative thought corresponds to reduced variability in the medial temporal system, and increased coupling between the frontoparietal network and default network core (2). The medial temporal lobe system therefore emerges as influential in the origin of spontaneous thoughts, with frontoparietal control regions becoming increasingly important for subsequent elaboration and metacognitive processing (6).

Exploring mind wandering in clinical populations can provide unique information about its cognitive and neural substrates. Altered mind wandering is documented in many conditions, and may constitute an important neurocognitive endophenotype across disorders. Perseverative mind wandering that is more frequent or salient, with negative content, has been reported in depressive rumination, neuroticism and dysphoria (7, 8), and is suggested to reflect an overly constrained mode of function in the default network, leading to excessive stability of thoughts (2). In contrast, higher rates of unintentional, spontaneous mind wandering are associated with increased obsessive-compulsive and attention-deficit/hyperactivity symptomatology in non-clinical samples (9, 10). Similarly, higher frequencies of mind wandering have been noted in schizophrenia, which correlate with the severity of positive symptoms (11). An unconstrained default network, due to local hyperactivity or relaxed influence from frontoparietal regions, may underpin excessive variation and incoherence of thoughts, as seen in psychosis (2).

Previous studies in neuropsychiatric populations have tended to explore network alterations with respect to specific maladaptive expressions of mind wandering, for example rumination in depression (12, 13). Notably, however, there exists a paucity of data directly relating brain network dysfunction to mind wandering capacity in clinical populations. As such, it remains unclear how pathological brain states impact the frequency and phenomenology of mind wandering.

The present study addresses this by directly testing whether pathological changes in the default and frontoparietal networks are associated with alterations in mind wandering capacity in neurodegenerative disorders. Dementia syndromes afford a unique opportunity to study the impact of network level dysfunction on mind wandering, given well established pathology primarily targeting, but not restricted to, nodes of the default and frontoparietal networks (14-16). This approach is an extension to recent work confirming that focal lesions to the default network, in the hippocampus and medial prefrontal cortex, can impact the content of mind wandering or reduce its frequency (17, 18). Moreover, on the cognitive level, many of the component processes implicated in mind wandering are disrupted in neurodegenerative disorders, for example autobiographical memory retrieval (19, 20), mental construction (21-23), working memory and shifting attention (24, 25).

Given these well-established neurocognitive changes, it follows that distinct alterations in the frequency and phenomenology of mind wandering should be present in dementia. A recent study reported increased on-task thoughts, suggestive of reduced mind wandering, during concurrent performance of a sustained attention task in mild Alzheimer’s disease (26). Reduced spontaneous mind wandering has been further been demonstrated in mild cognitive impairment during performance of a simple vigilance task (27). No study to date, however, has empirically investigated how alterations in structural and functional brain network integrity across dementia syndromes relate to mind wandering. This line of enquiry is crucial to establish how damage to brain networks impacts internally generated thought processes, whilst yielding important new insights into the cognitive symptomatology of these syndromes.

To this end, we explored mind wandering capacity in two dementia subtypes: Alzheimer’s disease (AD) and the behavioural variant of frontotemporal dementia (bvFTD). AD, characterised by prominent episodic memory deficits, is associated with pathological changes in the default and frontoparietal networks, particularly the hippocampus, medial temporal lobe subsystem and posterior cingulate cortex, extending into prefrontal and parietal regions with disease progression (15, 28, 29). In contrast, bvFTD is distinguished by behavioural dysfunction, including disinhibition, apathy, emotional blunting, stereotypical behaviours, and loss of insight. Early pathological changes in bvFTD target key regions of the salience and default networks, including the dorsomedial and ventromedial prefrontal cortices, as well as widespread changes across the amygdalae, thalamus and striatum with disease progression (15, 28, 30).

Given the marked deficits in cognitive mechanisms related to mind wandering in these two dementia syndromes, we predicted distinct changes in mind wandering profiles in both bvFTD and AD. We hypothesised that both groups would show an overall reduced propensity for mind wandering relative to controls. Given that environmentally dependent behaviours are characteristic of the bvFTD syndrome (31), we further predicted an increase in stimulus-bound forms of thought in bvFTD. Moreover, we sought to establish the neural correlates of mind wandering capacity in these syndromes, given their widespread structural and functional disruption to the default and frontoparietal networks.

Quantifying the nature and content of mind wandering in clinical disorders is inherently challenging. Dominant experimental approaches require subjects to monitor or self-identify extraneous thoughts during an ongoing cognitive task. Such approaches rely on dual-tasking and metacognitive capacities that are diminished in dementia, limiting the extent to which reliable conclusions can be drawn from existing measures. To circumvent these methodological constraints, we developed a paradigm to measure mind wandering under conditions of low cognitive demand (32). The task quantifies mind wandering as thoughts unrelated to the immediate environment or to the task at hand, consistent with current theoretical frameworks in which mind wandering is operationalised as stimulus-independent task-unrelated thought (33, 34). Thoughts are therefore classified along a conceptual continuum ranging from stimulus-bound, to stimulus-/task-related, through to fully-fledged instances of mind wandering (i.e., stimulus-independent task-unrelated thought).

The objectives of the current study were twofold. First, we aimed to quantify the capacity for mind wandering in dementia syndromes during conditions of low cognitive demand. Second, we sought to characterise how disease-related alterations in (i) regional grey matter, and (ii) seed-based functional connectivity in the default and frontoparietal networks, relate to mind wandering performance. In doing so, we aimed to validate current frameworks of mind wandering in a clinical model, by showing that the integrity of the default and frontoparietal networks is essential to support mind wandering capacity.

## Results

### Overall mind wandering performance

The task scoring system conceptualises mind wandering along a continuum, ranging from Level 1 (stimulus-bound thought) to Level 4 (mind wandering). Figure 1a displays the percentage of responses at each scoring level across the 9 trials. The main finding was that bvFTD patients displayed significantly increased stimulus-bound responses (Level 1) in the context of significantly decreased mind wandering responses (Level 4), relative to controls.

A group (bvFTD, AD, control) by Level (1,2,3,4) repeated measures ANOVA revealed no main effect of group on the task (*F*(2, 93) = 1.13, *p* = .328). A significant main effect of response level (*F*(3, 279) = 4.82, *p* < .01), was driven by a significant difference between percentage of responses at Level 3 vs. Level 1 (*p* < .001). No other significant differences across response levels were observed (*p* values > .07).

The group by response level interaction was significant (*F*(6, 276) = 7.42, *p* < .00001) and followed by tests of simple effects. Responses differed significantly between the groups at Level 1 [simple effect, *F*(2, 93) = 14.51, *p* < .00001], Level 2 [simple effect, *F*(2,93) = 7.64, *p* < .001], and at Level 4 [simple effect, *F*(2, 93) = 5.81, *p* < .01]; the groups did not differ at Level 3 [simple effect, *F*(2, 93) = .590, *p* = .557]. Sidak-corrected pairwise comparisons confirmed that bvFTD patients provided significantly more stimulus-bound Level 1 responses than AD (*p* < .05) and controls (*p* < .00001) (AD vs controls *p* = .116). In contrast, bvFTD Level 4 responses were significantly reduced relative to controls (*p* < .01) indicating a significant reduction in mind wandering (AD vs. controls *p* = .142; AD vs. bvFTD *p* = .670). BvFTD Level 2 responses were also significantly reduced compared to AD and controls (*p* values < .01) (AD vs controls *p* = .795). Finally, all groups were found to display higher average scores on longer-duration trials, suggesting an increased propensity for mind wandering with increasing stimulus duration (See SI Appendix, Fig. S1).

**Figure 1.**
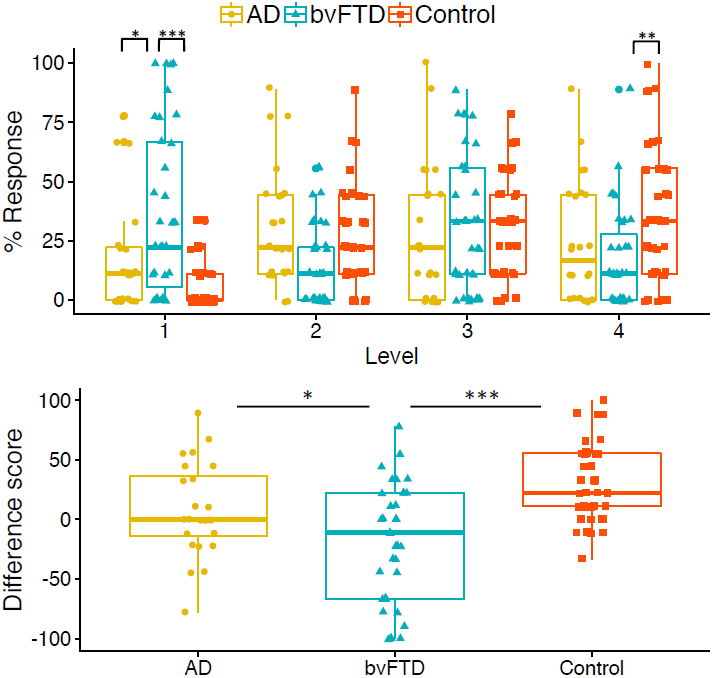
**a) Overall proportion of mind wandering scores across participant groups.** % responses across the mind wandering continuum. Asterisks show main results of group differences at Level 1 and Level 4. Level 1 responses represent stimulus-bound thoughts; Level 4 responses denote fully-fledged instances of mind wandering. **b) Average mind wandering index scores.** Mind wandering index (i.e., % difference in Level 4 minus Level 1 responses). Higher scores reflect an increased propensity to engage in mind wandering as opposed to stimulus-bound thought; with lower scores reflecting a tendency toward stimulus-bound thought. \**p* < .05; \*\**p* < .01; \*\*\**p* < .001.

To explore group differences in the overall pattern of responses we performed a linear trend analysis. This was to determine if observed responses across the levels were best described by a linear fit, such that participants would have a progressively higher percentage of responses across Levels 1-4 consistent with the response profile predicted for healthy controls (32). In line with our predictions, controls’ data was well fit by a linear model [*F*(1, 146) = 44.65, *p* < .00001, with an *R^2^* of 0.23]. In contrast, a significant linear trend was not observed in either the bvFTD [*F*(1,138) = 2.28, *p* = 0.13, with an *R^2^* of 0.02] or the AD [*F*(1, 94) = .53, *p* = .47, with an *R^2^* of 0.006] group, suggesting that the overall response profile in the two dementia groups differed from that of controls.

### Mind wandering index score

To compare the proportion of Level 1 (stimulus-bound) with Level 4 (mind wandering) responses, an index score was created by subtracting the % of Level 1 responses from % Level 4. A larger positive index score reflects a tendency to engage in mind wandering as opposed to stimulus-bound thought, with negative scores reflecting the reverse profile. Figure 1b shows the average mind wandering index score across participant groups. Significant group differences were observed on the mind wandering index (*F*(2, 93) = 13.57, *p* < .00001). Sidak-corrected pairwise comparisons confirmed the bvFTD group scored significantly lower than both AD (*p* < .05) and controls (*p* < .00001), while AD patients did not differ significantly from controls (*p* = .110).

### Grey matter correlates of mind wandering performance

Voxel-based morphometry was used to determine the relationship between the mind wandering index and regional grey matter intensity in the patient groups. Figure 2 displays clusters in which a significant positive correlation emerged between grey matter intensity and the mind wandering index in both bvFTD and AD groups combined. Reduced grey matter intensity in these regions was associated with lower mind wandering index scores, reflecting the tendency towards stimulus-bound thought. Three main clusters were identified: i) striatum (including caudate, putamen, nucleus accumbens) and anterior-mid thalamus, extending to the left subcallosal, medial/lateral orbitofrontal, and anterior insular cortices; ii) left hippocampus and parahippocampal gyrus; iii) left posterior insular cortex (See SI Appendix, Table S2 for co-ordinates). These regions showed considerable overlap with areas of grey matter intensity reduction in the patient groups relative to controls (see SI Appendix, Table S4 and Fig. S3). To illustrate the resting state networks that these regions overlapped with, the results are overlaid on the Yeo et al. (35) 17-network cortical parcellation scheme (see Figure S4). Of note, the anterior insula cluster overlapped with the salience and default networks, the subcallosal/orbitofrontal cluster with the limbic network, and the parahippocampal cluster with the limbic and default networks.

**Figure 2.**
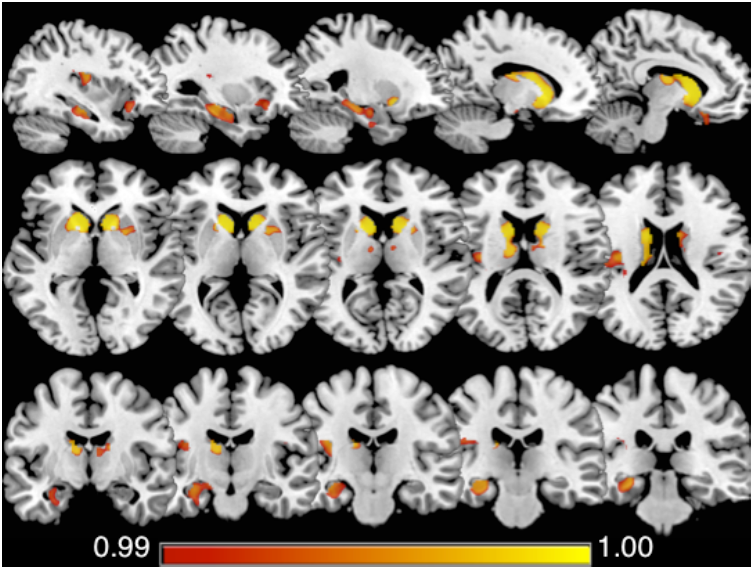
Regions of grey matter intensity that covaried with the mind wandering index in bvFTD and AD patients combined. Significant clusters were identified in the striatum (including caudate, putamen, nucleus accumbens) and anterior-mid thalamus, extending to the left subcallosal, medial/lateral orbitofrontal, and anterior insular cortices; the left hippocampus and parahippocampal gyrus; and the left posterior insular cortex. Results FWE corrected at *p* < .01; significant clusters identified using threshold free cluster enhancement.

### Seed region connectivity and mind wandering performance

The relationship between seed region connectivity and the mind wandering index score was examined using seeds placed within the default and frontoparietal networks, and hippocampus. Connections that significantly correlated with the mind wandering index within each group are shown in Figure 3. In controls, the tendency to mind wander (as opposed to stimulus-bound thought) was positively associated with left PCC-left posterior hippocampus connectivity (*r* = .14, *q* < .05) and negatively associated with left dlPFC-left hippocampal formation connectivity (*r* = -.23, *q* < .05) and left PCC-vmPFC (*r* = -.30, *q* < .05) connectivity.

The AD group showed the opposite pattern for left PCC-left posterior hippocampus connectivity, as this was negatively associated with a tendency to mind wander (*r* = −.56, *q* < .05; this was significantly different to controls’ connectivity between the same regions: Z = −2.09, *p* < .05). The tendency to mind wander in the AD group was also negatively associated with left hippocampal formation-vmPFC connectivity (*r* = -.62, *q* < .05).

Similar to the AD group, bvFTD patients showed the opposite (negative) relationship to controls for left PCC-left posterior hippocampus connectivity (*r* = −.44, *q* < .05; which differed significantly from controls’ connectivity between the same regions: Z = −1.91, *p* < .05, but not from ADs’ connectivity between those regions: Z = −.43, *p* = .33). The bvFTD group also showed the opposite relationship to controls for left dlPFC-left hippocampal formation connectivity and left PCC-vmPFC connectivity, which were both positively associated with a tendency to mind wander (dlPFC-HF: *r* = .48, *q* < .05; PCC-vmPFC: *r* = .58, *q* < .05) and both of which were significantly different to controls (dlPFC-HF: Z = 2.30, *p* < .05; PCC-vmPFC: Z = 2.97, *p* < 0.01). bvFTD showed additional significant positive associations with mind wandering, between left dlPFC-left hippocampal formation connectivity (*r* = .48, *q* < .05) and right PCC-right-amPFC connectivity (*r* = .44, *q* < .05), as well as significant negative associations with left dlPFC-right amPFC connectivity (*r* = −.47, *q* < .05), right PCC-left posterior hippocampal connectivity (*r* = −.51, *q* < .05), and right posterior hippocampus-right hippocampal formation connectivity (*r* = −.46, *q* < .05).

**Figure 3.**
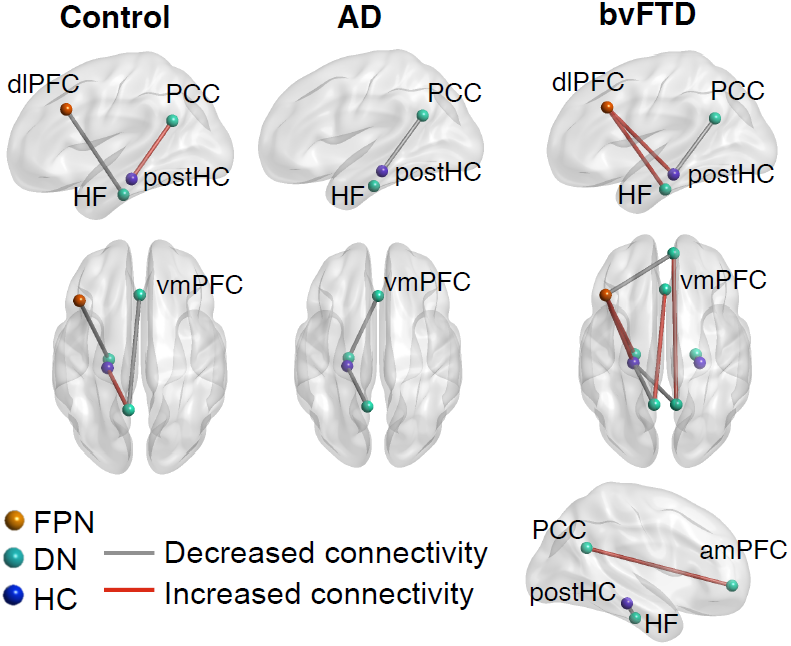
Seed regions where connectivity was associated with a tendency to mind wander within the participant groups. Connections show where the mind wandering index was significantly correlated with connectivity, such that a tendency to mind wander on the task (as opposed to stimulus-bound thought) was either positively (red lines) or negatively (grey lines) associated with connectivity. Colour of ROIs correspond to the networks the regions are taken from: FPN = frontoparietal network; DN = default network; HC = hippocampus. Results are FDR corrected at *q* < .05. dlPFC = dorsolateral prefrontal cortex; PCC = posterior cingulate cortex; postHC = posterior hippocampus; HF = hippocampal formation; vmPFC = ventromedial prefrontal cortex; amPFC = anteromedial prefrontal cortex. Left view = left side of brain; Right view = right side of brain.

## Discussion

This study offers some of the first insights into mind wandering capacity, and its associated neural substrates, across dementia syndromes. Our results point to shifts in the thought profiles elicited during periods of low cognitive demand in dementia, which in turn are associated with structural integrity and resting-state functional connectivity in the default and frontoparietal networks. These findings corroborate current theoretical frameworks emphasising the role of the medial temporal lobe, and interactions between the default and frontoparietal networks, in supporting stimulus-independent task-unrelated thought (2).

Our most striking behavioural finding was a bias toward stimulus-bound thought in bvFTD. Relative to controls, the bvFTD group displayed significantly more instances of stimulus-bound thought (Level 1) in the context of significantly reduced mind wandering (Level 4). This pattern was reflected by a lower, negative mind wandering index in bvFTD relative to the other groups. By contrast, healthy older controls displayed a larger, positive mind wandering index, reflecting preferential engagement in perceptually decoupled thought. Our findings suggest that bvFTD patients experience marked difficulties in disengaging from the immediate environment, leading to a predominantly stimulus-bound style of thought.

In contrast, no clear differences emerged between the AD group and controls across any of the response levels or the mind wandering index. However, the distribution of responses across the four levels in AD suggested subtle alterations in their overall response profile. Controls’ responses showed a positive linear trend across the four levels, consistent with a greater proportion of responses at higher levels on the task, replicating previous findings (32). This trend was not observed in AD, with equivalent frequency of thoughts at each response level suggesting a shift from the normative profile. We tentatively suggest that in early stages of AD, thought content gradually shifts towards intermediate and increasingly stimulus-related forms of thought, leading to a comparable distribution of responses across task levels. Importantly, the disproportionate impairment in mind wandering in bvFTD relative to AD cannot be attributed to greater disease severity in the bvFTD group. The patient groups were matched for overall disease duration, with characteristic profiles of greater behavioural impairment in bvFTD and greater overall cognitive impairment in AD (cf. table S1 results of the cognitive and behavioural screening assessments). Rather, our results suggest that under conditions of low cognitive demand, AD patients can achieve a form of mind wandering despite their underlying widespread structural and functional brain changes.

Research investigating mind wandering in dementia is scarce, however a recent study revealed increased on-task thoughts during performance of an ongoing task (i.e., the SART), in early AD relative to controls (26). This finding differs from the current results, where we did not see an obvious reduction in mind wandering in AD. It is important to note, however, that the paradigms used in the two studies differ considerably in the cognitive demands imposed. Performance of an ongoing task is inherently more cognitively demanding than the thought sampling paradigm used in the current study. For patients with AD, the increased attentional demands of the SART invariably leave less cognitive resources available, reducing the likelihood of engaging in mind wandering (26). In contrast, the very low cognitive demands of our task render it more conducive to mind wandering in the AD group. These differences across studies emphasise that dual-tasking requirements may be a crucial determinant of ongoing thought patterns in cognitively impaired groups. We note with interest that when cognitive demands are lessened, AD patients appear capable of engaging in task-independent forms of thought. Whether the content, phenomenology, and intentional versus unintentional nature of mind wandering in AD differs from that of controls remains an important question for future study. Consistent with this possibility, individuals with selective bilateral hippocampal damage have been shown to mind wander as frequently as controls, albeit with distinct differences in terms of content (18).

Substantial evidence has shown that component processes supporting mind wandering, particularly those involving memory-based constructive simulation, are compromised in dementia (36, 37). AD and bvFTD patients display comparable episodic memory dysfunction (38-40). Both groups also display marked impairments in future-oriented thinking, including prospective memory (41, 42) and constructive simulation of future episodes (21, 43, 44). Accordingly, similar reductions in mind wandering might have been predicted. Our results, however, suggest that mind wandering capacity is more vulnerable in bvFTD, consistent with other features of the syndrome that point to a predisposition for stimulus-bound thought. The stimulus-bound thought style we observed in bvFTD resonates with reports of environmentally dependent behaviour in their everyday life, including preoccupation with objects in the immediate environment and the inability to disengage from such external stimuli (45, 46). We suggest that the current findings capture a core feature of the broader bvFTD behavioural phenotype, not previously reported, namely a change in spontaneous thought style. Naturalistic paradigms assessing mind wandering over extended time periods (e.g., 18) would be an important extension to the current study, and would enable us to determine how task demands and contextual factors influence stimulus-bound thought in bvFTD.

Our neuroimaging findings underscore the role of key regions of the default and frontoparietal networks in supporting internally generated thought, corroborating previous reports in healthy individuals (3-5). The resting state results showed that, in controls, the tendency to mind wander (as opposed to stimulus-bound thought) was associated with stronger left PCC-left posterior hippocampal connectivity, and weaker left PCC-vmPFC and left dlPFC-left hippocampal formation connectivity. This suggests that relative engagement of a posterior memory system and disengagement of frontal systems may be important for mind wandering on our task, in healthy individuals.

In the patient groups, we observed the opposite association between left PCC-left posterior hippocampal connectivity, as mind wandering was associated with weaker connectivity between these regions. The bvFTD group showed additional opposite associations to controls, with stronger left PCC-vmPFC and left dlPFC-left hippocampal formation connectivity associated with mind wandering. This suggests that for bvFTD, mind wandering in the context of our task may rely more on frontal systems. In general, bvFTD showed more widespread associations between mind wandering and connectivity, relative to both controls and AD. This pattern may reflect a compensatory re-organisation of networks involved in mind wandering, or it may reflect dedifferentiation, that is, a loss of specialisation in the regions or systems supporting a given function. Both of these mechanisms can occur in ageing and neurodegeneration in response to local atrophy or structural change (47-49), and future work is needed to disambiguate between these processes in the context of mind wandering in bvFTD. Our results suggest that in bvFTD, and to a lesser extent in AD, perceptually-decoupled thought may be supported by a different functional architecture or by different psychological characteristics than controls, which in turn may contribute to the observed behavioural differences.

In both patient groups, a tendency toward stimulus-bound thought at the expense of mind wandering (i.e., lower mind wandering index), was associated with decreased grey matter intensity in several regions. Most notable in the context of existing literature were associations between the mind wandering index and grey matter integrity in the left hippocampus and parahippocampus. Convergent measures show that activity in the hippocampus and parahippocampus (6), entorhinal cortex (50) and temporal cortex (51) precedes spontaneous free recall of episodic memories. This accords with rodent work implicating hippocampal sharp wave ripple events (SWRs) in the replay (and preplay) of previously learnt or future behavioural sequences (52, 53), and in monkeys, where hippocampal SWRs precede increased activation of the default network (54). Together, these findings link spontaneous activation in the hippocampus and surrounding regions with both recall and prospection, which may then engage the default network more broadly to support the elaboration of memories and simulations. A critical role for the hippocampus in mind wandering was also recently confirmed, as individuals with selective bilateral hippocampal damage exhibited reduced diversity in their mind wandering content (18). We extend these findings by demonstrating that decreased grey matter intensity in the hippocampus/parahippocampus is associated with an increased propensity to engage in stimulus-bound thought during periods of low cognitive demand.

Significant associations were also observed between the mind wandering index score and grey matter integrity in areas overlapping with large-scale networks relevant for mind wandering. Grey matter clusters in the subcallosal/orbitofrontal and parahippocampal cortices overlapped with the limbic and default networks. Furthermore, a cluster in the anterior insula overlapped with the salience network. The salience network is proposed to mediate dynamic shifts between default and executive control networks (55) facilitating transitions between external and internal focus, which may be relevant for disengaging from external stimuli to engage in internally focused thought. While the structural correlates of task-independent thought have received less attention relative to the functional correlates, cortical thickness in regions within and adjacent to the default and frontoparietal networks has been shown to covary with mind wandering performance in healthy individuals (56). The implication is that structural integrity in regions within and adjacent to these networks helps to promote their integration (56).

Finally, we found an association between striatal grey matter loss and a reduced mind wandering index. The basal ganglia represents a network hub vulnerable to degeneration across neurodegenerative disorders (57), contributing to an array of cognitive and neuropsychiatric features (58). We speculate that the involvement of the basal ganglia in supporting large scale network communication (59) may explain its association with mind wandering. This is consistent with known functional connectivity between the striatum and large-scale cortical networks, including default and frontoparietal (60), and the convergence of these functional networks in distinct zones of the striatum (61). Striatal degeneration may impair the integration of information from disparate brain networks, which is necessary to support abstract forms of cognition, including mind wandering. As the dynamic integration and segregation of network communication is increasingly recognised as important for mind wandering (1, 62), further investigation of the striatal influence over large scale network topology in this context is warranted.

In summary, to our knowledge, this is the first study to empirically measure mind wandering under conditions of low cognitive demand in two dementia syndromes, and to reconcile performance with structural and functional imaging correlates. Our results show a change in the thought patterns of individuals with bvFTD, and to a much lesser extent those with AD. The tendency to engage in mind wandering vs. stimulus bound thought was associated with regional grey matter integrity, and functional connectivity in the default and frontoparietal networks. Future work is needed to identify the trait level and phenomenological characteristics of altered mind wandering in dementia. Given the ubiquity of mind wandering in everyday life, we also stress the importance of further understanding how the loss of this fundamental human capacity impacts wellbeing and sense of self in individuals living with dementia.

## Methods and Materials

### Case selection

The study included 35 individuals meeting diagnostic criteria for bvFTD, 24 individuals with a clinically probable diagnosis of AD, and 37 healthy controls. See SI Appendix, Materials and Methods and Table S1. The South Eastern Sydney Local Area Health and University of New South Wales ethics committees approved the study and all participants provided informed consent. The data supporting this study is unavailable as ethics did not cover open data sharing, however stimulus materials for the task are available from the authors upon request.

### Mind wandering experimental task

Participants viewed static, two-dimensional coloured geometric shapes presented individually on a computer screen. Immediately following the presentation of each stimulus, participants were prompted to report their thoughts that arose during stimulus presentation. The task comprised 9 trials, each presenting a commonplace shape (e.g., blue square, yellow circle) for varying durations (Short: ≤20 s, Medium: 30-60 s, Long: ≥90 s). The scoring procedure for this task has been described previously (32, 63). Briefly, responses are coded along a continuum ranging from stimulus-bound thinking directly related to the stimulus at hand (Level 1) to fully-fledged instances of stimulus-/task-unrelated mind wandering (Level 4). Intermediary levels 2 and 3 capture the transition from stimulus-related to increasingly stimulus-independent responses. The final score awarded for each trial was the highest level achieved on that trial, ranging from 1-4. Total percentages of each level across the task were calculated, as well as the mind wandering index comparing the extent of Level 1 vs. Level 4 responses. The mind wandering index was used as the covariate of interest in the neuroimaging, rather than mind wandering percentage (Level 4) as many patients scored 0 for this, leading to reduced variance in the sample. See SI Appendix for the scoring protocol and representative responses.

### VBM analysis of mind wandering performance

Structural scans were available for 31 bvFTD, 23 AD, and 32 controls. To identify regions where grey matter intensity covaried with mind wandering performance, a GLM was conducted in the patient groups combined (i.e., excluding controls), using the mind wandering index score as a covariate in the design matrix. A priori the regions of interest were specified as all cortical and subcortical regions within the Harvard-Oxford cortical and subcortical atlases; the cerebellum and brain stem were not included. We tested for interaction effects, and having confirmed that they were not significant, the main effect of mind wandering index on grey matter intensity was reported, i.e., regions where a significant positive relationship with the mind wandering index in both groups combined was identified. Results are FWE corrected at p < .01 and clusters identified using threshold free cluster enhancement. (See SI Appendix for details, pre-processing procedures and group-level comparisons).

### Seed region connectivity and mind wandering performance

A subset of 24 bvFTD, 17 AD, and 23 controls underwent task-free resting state imaging (3 bvFTD and 3 AD were removed from the analysis due to excessive motion; see SI Appendix for details). Thirteen seeds were placed in the default and frontoparietal networks, and hippocampus, to determine the relationship between seed region connectivity and the mind wandering index. Within each of the three groups, participants’ mind wandering index scores were correlated with the connectivity between each ROI for the 13 seed-regions of interest. Correlations that survived FDR correction at *q* < 0.05 are reported. To compare the strength of shared correlations across groups, Fisher’s r to z was calculated and a one-tailed comparison at *p* < .05 (i.e. Z ≥ ± 1.645) is reported. (See SI Appendix for details, pre-processing procedures and group-level comparisons.)

## Acknowledgments

We thank Nadene Dermody and Jody Kamminga for assistance with data collection and scoring. CO is supported by a National Health and Medical Research Council (NHMRC) Neil Hamilton Fairley Fellowship GNT1091310 and by the Wellcome Trust (200181/Z/15/Z). JMS is supported by a NHMRC CJ Martin Fellowship GNT1072403. JAH is supported by the University of Arizona and a grant from the John Templeton Foundation, “Prospective Psychology Stage 2: A Research Competition” to Martin Seligman. The opinions expressed in this publication are those of the author(s) and do not necessarily reflect the views of the John Templeton Foundation. MI is supported by an Australian Research Council (ARC) Future Fellowship (FT160100096) and an ARC Discovery Project (DP180101548). This work is supported by funding to ForeFront, a collaborative research group dedicated to the study of frontotemporal dementia and motor neuron disease, from the NHMRC (APP1037746) and the ARC Centre of Excellence in Cognition and its Disorders (CE11000102).

## References

1. Kucyi A (2017) Just a thought: How mind-wandering is represented in dynamic brain connectivity. NeuroImage.

2. Christoff K, Irving ZC, Fox KC, Spreng RN, & Andrews-Hanna JR (2016) Mind-wandering as spontaneous thought: a dynamic framework. Nature Reviews Neuroscience 17(11):718–731.

3. Christoff K, Gordon AM, Smallwood J, Smith R, & Schooler JW (2009) Experience sampling during fMRI reveals default network and executive system contributions to mind wandering. Proceedings of the National Academy of Sciences 106(21):8719–8724.

4. Fox KC, Spreng RN, Ellamil M, Andrews-Hanna JR, & Christoff K (2015) The wandering brain: Meta-analysis of functional neuroimaging studies of mind-wandering and related spontaneous thought processes. NeuroImage 111:611–621.

5. Mason MF, et al. (2007) Wandering minds: the default network and stimulus-independent thought. Science 315(5810):393–395.

6. Ellamil M, et al. (2016) Dynamics of neural recruitment surrounding the spontaneous arising of thoughts in experienced mindfulness practitioners. NeuroImage 136:186–196.

7. Smallwood J, O'Connor RC, Sudbery MV, & Obonsawin M (2007) Mind-wandering and dysphoria. Cognition and Emotion 21(4):816–842.

8. Perkins AM, Arnone D, Smallwood J, & Mobbs D (2015) Thinking too much: self-generated thought as the engine of neuroticism. Trends in cognitive sciences 19(9):492–498.

9. Seli P, Smallwood J, Cheyne JA, & Smilek D (2015) On the relation of mind wandering and ADHD symptomatology. Psychonomic bulletin & review 22(3):629–636.

10. Seli P, Risko EF, Purdon C, & Smilek D (2016) Intrusive thoughts: linking spontaneous mind wandering and OCD symptomatology. Psychological research:1–7.

11. Shin D-J, et al. (2015) Away from home: the brain of the wandering mind as a model for schizophrenia. Schizophrenia research 165(1):83–89.

12. Berman MG, et al. (2014) Does resting-state connectivity reflect depressive rumination? A tale of two analyses. Neuroimage 103:267–279.

13. Hamilton JP, et al. (2011) Default-Mode and Task-Positive Network Activity in Major Depressive Disorder: Implications for Adaptive and Maladaptive Rumination. Biological Psychiatry 70(4):327–333.

14. Irish M, Piguet O, & Hodges JR (2012) Self-projection and the default network in frontotemporal dementia. Nature Reviews Neurology 8(3):152–161.

15. Zhou J, et al. (2010) Divergent network connectivity changes in behavioural variant frontotemporal dementia and Alzheimer’s disease. Brain 133(5):1352–1367.

16. Pievani M, de Haan W, Wu T, Seeley WW, & Frisoni GB (2011) Functional network disruption in the degenerative dementias. The Lancet Neurology 10(9):829–843.

17. Bertossi E & Ciaramelli E (2016) Ventromedial prefrontal damage reduces mind-wandering and biases its temporal focus. Social Cognitive and Affective Neuroscience 11(11):1783–1791.

18. McCormick C, Rosenthal CR, Miller TD, & Maguire EA (2018) Mind-wandering in people with hippocampal damage. Journal of Neuroscience:1812–1817.

19. Irish M, et al. (2011) Profiles of recent autobiographical memory retrieval in semantic dementia, behavioural-variant frontotemporal dementia, and Alzheimer's disease. Neuropsychologia 49(9):2694–2702.

20. Piolino P, et al. (2003) Autobiographical memory and autonoetic consciousness: triple dissociation in neurodegenerative diseases. Brain 126(10):2203–2219.

21. Irish M, et al. (2015) Scene construction impairments in Alzheimer's disease–A unique role for the posterior cingulate cortex. cortex 73:10–23.

22. Irish M, Addis DR, Hodges JR, & Piguet O (2012) Considering the role of semantic memory in episodic future thinking: evidence from semantic dementia. Brain 135(7):2178–2191.

23. Duval C, et al. (2012) What happens to personal identity when semantic knowledge degrades? A study of the self and autobiographical memory in semantic dementia. Neuropsychologia 50(2):254–265.

24. Stopford CL, Thompson JC, Neary D, Richardson AMT, & Snowden JS (2012) Working memory, attention, and executive function in Alzheimer’s disease and frontotemporal dementia. Cortex 48(4):429–446.

25. Possin KL, et al. (2013) Dissociable executive functions in behavioral variant frontotemporal and Alzheimer dementias. Neurology 80(24):2180–2185.

26. Gyurkovics M, Balota D, & Jackson J (2018) Mind-Wandering in Healthy Aging and Early Stage Alzheimer's Disease. Neuropsychology 32(1):89–101.

27. Niedzwienska A & Kvavilashvili L (2018) Reduced mind-wandering in mild cognitive impairment: Testing the spontaneous retrieval deficit hypothesis. Neuropsychology.

28. Seeley WW, Crawford RK, Zhou J, Miller BL, & Greicius MD (2009) Neurodegenerative diseases target large-scale human brain networks. Neuron 62(1):42.

29. Braak H & Braak E (1991) Neuropathological stageing of Alzheimer-related changes. Acta neuropathologica 82(4):239–259.

30. Broe M, et al. (2003) Staging disease severity in pathologically confirmed cases of frontotemporal dementia. Neurology 60(6):1005–1011.

31. Snowden J, et al. (2001) Distinct behavioural profiles in frontotemporal dementia and semantic dementia. Journal of Neurology, Neurosurgery & Psychiatry 70(3):323–332.

32. O’Callaghan C, Shine JM, Lewis SJG, Andrews-Hanna JR, & Irish M (2015) Shaped by our thoughts – A new task to assess spontaneous cognition and its associated neural correlates in the default network. Brain and Cognition 93(0):1–10.

33. Seli P, et al. (2018) Mind-wandering as a natural kind: A family-resemblances view. Trends in Cognitive Sciences.

34. Smallwood J & Schooler JW (2015) The science of mind wandering: empirically navigating the stream of consciousness. Annual review of psychology 66:487–518.

35. Yeo BT, et al. (2011) The organization of the human cerebral cortex estimated by intrinsic functional connectivity. Journal of Neurophysiology 106(3):1125–1165.

36. Irish M & Piolino P (2016) Impaired capacity for prospection in the dementias– Theoretical and clinical implications. British Journal of Clinical Psychology 55(1):49–68.

37. O’Callaghan C & Irish M (2018) Candidate mechanisms of spontaneous cognition as revealed by dementia syndromes. The Oxford Handbook of Spontaneous Thought, eds Fox KCR & Christoff K (Oxford University Press, New York).

38. Bertoux M, et al. (2016) Social cognition deficits: the key to discriminate behavioral variant frontotemporal dementia from Alzheimer’s disease regardless of amnesia? Journal of Alzheimer’s Disease 49(4):1065–1074.

39. Hornberger M, Piguet O, Graham A, J., Nestor PJ, & Hodges JR (2010) How preserved is episodic memory in behavioral variant frontotemporal dementia? Neurology 74:472–479.

40. Irish M, Piguet O, Hodges JR, & Hornberger M (2014) Common and unique gray matter correlates of episodic memory dysfunction in frontotemporal dementia and alzheimer’s disease. Human Brain Mapping 35(4):1422–1435.

41. Kamminga J, O’Callaghan C, Hodges JR, & Irish M (2014) Differential Prospective Memory Profiles in Frontotemporal Dementia Syndromes. Journal of Alzheimer’s Disease 38(3):669–679.

42. Dermody N, Hornberger M, Piguet O, Hodges JR, & Irish M (2016) Prospective memory impairments in Alzheimer’s disease and behavioral variant frontotemporal dementia: Clinical and neural correlates. Journal of Alzheimer’s Disease 50(2):425–441.

43. Addis DR, Sacchetti DC, Ally BA, Budson AE, & Schacter DL (2009) Episodic simulation of future events is impaired in mild Alzheimer’s disease. Neuropsychologia 47(12):2660–2671.

44. Irish M, Hodges JR, & Piguet O (2013) Episodic future thinking is impaired in the behavioural variant of frontotemporal dementia. Cortex 49(9):2377–2388.

45. Lanata SC & Miller BL (2016) The behavioural variant frontotemporal dementia (bvFTD) syndrome in psychiatry. J Neurol Neurosurg Psychiatry 87(5):501–511.

46. Ghosh A, Dutt A, Bhargava P, & Snowden J (2013) Environmental dependency behaviours in frontotemporal dementia: have we been underrating them? Journal of neurology 260(3):861–868.

47. Kalpouzos G, Persson J, & Nyberg L (2012) Local brain atrophy accounts for functional activity differences in normal aging. Neurobiology of Aging 33(3):623.e621–623.e613.

48. Maillet D & Rajah MN (2013) Association between prefrontal activity and volume change in prefrontal and medial temporal lobes in aging and dementia: A review. Ageing Research Reviews 12(2):479–489.

49. O’Callaghan C, et al. (2016) Cerebellar atrophy in Parkinson’s disease and its implication for network connectivity. Brain 139(3):845–855.

50. Gelbard-Sagiv H, Mukamel R, Harel M, Malach R, & Fried I (2008) Internally generated reactivation of single neurons in human hippocampus during free recall. Science 322(5898):96–101.

51. Burke JF, et al. (2014) Theta and high-frequency activity mark spontaneous recall of episodic memories. Journal of Neuroscience 34(34):11355–11365.

52. Pfeiffer BE & Foster DJ (2013) Hippocampal place-cell sequences depict future paths to remembered goals. Nature 497(7447):74–79.

53. Diba K & Buzsáki G (2007) Forward and reverse hippocampal place-cell sequences during ripples. Nature neuroscience 10(10):1241–1242.

54. Kaplan R, et al. (2016) Hippocampal sharp-wave ripples influence selective activation of the default mode network. Current Biology 26(5):686–691.

55. Menon V & Uddin LQ (2010) Saliency, switching, attention and control: a network model of insula function. Brain Structure and Function 214(5-6):655–667.

56. Golchert J, et al. (2017) Individual variation in intentionality in the mind-wandering state is reflected in the integration of the default-mode, fronto-parietal, and limbic networks. NeuroImage 146:226–235.

57. Crossley NA, et al. (2014) The hubs of the human connectome are generally implicated in the anatomy of brain disorders. Brain 137(8):2382–2395.

58. O’Callaghan C, Bertoux M, & Hornberger M (2014) Beyond and below the cortex: the contribution of striatal dysfunction to cognition and behaviour in neurodegeneration. Journal of Neurology, Neurosurgery and Psychiatry 85(4):371–378.

59. Bell PT & Shine JM (2016) Subcortical contributions to large-scale network communication. Neuroscience & Biobehavioral Reviews 71:313–322.

60. Choi EY, Yeo BT, & Buckner RL (2012) The organization of the human striatum estimated by intrinsic functional connectivity. Journal of neurophysiology 108(8):2242–2263.

61. Jarbo K & Verstynen TD (2015) Converging structural and functional connectivity of orbitofrontal, dorsolateral prefrontal, and posterior parietal cortex in the human striatum. Journal of Neuroscience 35(9):3865–3878.

62. Zabelina DL & Andrews-Hanna JR (2016) Dynamic network interactions supporting internally-oriented cognition. Current Opinion in Neurobiology 40:86–93.

63. Irish M, Goldberg Z-l, Alaeddin S, O’Callaghan C, & Andrews-Hanna JR (2018) Age-related changes in the temporal focus and self-referential content of spontaneous cognition during periods of low cognitive demand. Psychological research:1–14.

